# Late induction of IgG4 following SARS-CoV-2 mRNA vaccination in pregnant and non-pregnant individuals includes clonotypes raised early in the response

**DOI:** 10.64898/2026.05.29.728639

**Authors:** Dhvanir N. Kansara, Tae-Geun Yu, Neel Kansara, Noor M. Taher, Aisha Yesbolatova, Larissa DeBrabandere, Joshua A. Weiner, Gabriela Kovacikova, Andrew P. Hederman, Pieter Pannus, Stéphanie Depickère, Maria E. Goossens, An Vercoutere, Nicolas Dauby, Kevin K. Ariën, Kirsten Maertens, Arnaud Marchant, Jiwon Lee, Margaret E. Ackerman

## Abstract

To better understand how pregnancy impacts humoral immunity, we conducted an in-depth longitudinal analysis of the kinetics and characteristics of vaccine responses in a prospective cohort of pregnant and non-pregnant women. Humoral immune responses observed among pregnant participants who received the mRNA-delivered SARS-CoV-2 vaccination, including their effector functions, were in some cases marginally lower than those among non-pregnant controls, while prior infection was associated with some potentiation in humoral responses. Importantly, vaccine-induced antibodies were efficiently transferred across the placenta, providing the fetus with passive immunity and underscoring the dual benefit of maternal vaccination for both mother and neonate against COVID-19. Delayed induction of spike-specific IgG4 following the primary two-dose vaccination series was observed in vaccine recipients, independent of pregnancy status. In a subset (n=6) of pregnant women whose spike-specific serum IgG repertoires were extensively profiled at the clonotypic level over time as part of another study, we proteomically identified secreted IgG clonotypes that had class-switched to IgG4. Matching of these clonotypes detected as IgG4 to those defined as SARS-CoV-2 spike-specific revealed that, while a minority of total clonotypes, they were elicited early in the immunization series and tended to be more highly mutated, more prevalent, and more persistent than clonotypes in the serological repertoire that were not detected as IgG4. Consistent with the increase in secreted vaccine-specific IgG4 over time, but its poorer placental transfer, these clonotypes were detected at greater levels in maternal but not cord blood at the time of delivery as compared to 28 days post the second vaccine dose. These findings indicate some impact in the kinetics, characteristics, and functions of the humoral response that may be associated with pregnancy-related immune modulation. Conservation of the late class-switch recombination to IgG4 that has previously been associated with mRNA-based SARS-CoV-2 vaccines raises questions about how different immunological states and vaccine components influence short- and long-term characteristics of the humoral immune response.

## Introduction

Especially early in the SARS-CoV-2 pandemic when most individuals were immunologically naive, pregnant women were at higher risk for severe disease and complications from infection, including hospitalization, requirement for respiratory support, intensive care unit admission, and, in some cases, mortality, compared to non-pregnant adults ^1-9^. Beyond maternal morbidity, infection during pregnancy has been associated with SARS-CoV-2 placentitis, which has been described even in the absence of severe maternal disease, and may result in placental ischemia, thereby increasing the risk of fetal growth problems, stillbirth, and preeclampsia ^5,10-12^. However, when the first COVID-19 vaccines became available, a significant gap in knowledge regarding their safety in pregnant women remained, as most Phase III clinical trials did not include this population ^13,14^. Fortunately, reassuring safety data have since accumulated, and robust antibody responses following vaccination in this population have been demonstrated, resulting in reduced risk of severe symptoms, complications, and death from COVID-19 when women are vaccinated during pregnancy ^15-22^. Hence, while vaccination is recommended in the general population, it is particularly important for pregnant women to reduce COVID-associated complications and maternal and infant morbidity and mortality.

Nonetheless, differences in immune responses associated with pregnancy may persist. Evidence regarding whether vaccine-induced humoral immune responses in pregnant women differ from those in non-pregnant counterparts remains equivocal for SARS-CoV as well as other vaccines ^23^. Overall, current immunogenicity data do not establish that pregnancy is associated with attenuated antibody responses, including diminished seroconversion rates, geometric mean titers (GMTs), or neutralizing antibody levels following immunization. Although pregnancy is characterized by well-documented trimester-dependent immunomodulatory adaptations ^24-26^, the precise impact of these physiological alterations on vaccine-elicited immunogenicity remains incompletely elucidated ^27-29^. Furthermore, considerable heterogeneity across study populations, vaccine platforms, timing of immunization relative to gestational age, and variability in serological assay methodologies may substantially contribute to the inconsistent and sometimes contradictory reports. Potential differences notwithstanding, data from the CDC indicate that maternal mRNA vaccination is not only effective in mothers, but 61% effective in preventing COVID-19-related hospitalization in newborns during their first six months of life ^30^. Thus, whether there are or are not generalizable differences in vaccine responses and resulting immunogenicity during pregnancy, maternal vaccination not only offers protection to the pregnant vaccinee but also provides inherited benefits to the fetus and neonate by reducing the risk of severe maternal illness and facilitating the transfer of COVID-specific IgG antibodies via the placenta and IgA through breast milk to protect infants post-partum ^31^.

Beyond their neutralization capacities, antibodies confer protection against COVID-19 by engaging the innate immune system through their Fc domains, potentially providing benefits that can extend to neutralization-resistant variants ^32,33^. Fc-mediated effector functions involve recruitment and activation of innate immune cells and the complement cascade, which operate independently of, or synergistically with, Fab domain-mediated neutralization capability ^34^. Although mRNA vaccines have been shown to induce robust humoral responses in pregnant and lactating women ^35^, prior studies on SARS-CoV-2 ^36-41^ have shown that cord blood levels of functional antibodies do not precisely reflect maternal antibody levels. Qualitative and quantitative differences in the transferred antibodies have been attributed to factors including placental factors and maternal antibody levels, glycosylation patterns, antigen specificity, and IgG subclass ^42-45^. Among these factors and across vaccines, differences in the levels of IgG1, IgG2, IgG3, and IgG4 in maternal versus cord blood are consistent observations ^46^. Given the striking functional differences between human IgG subclass, transfer efficiency differences may be of importance to robust protection of neonates.

Subclass-associated differences are of particular interest in the context of SARS-CoV-2 immunization during pregnancy. Firstly, prior to widespread seroconversion across the population, vaccination during pregnancy typically induced the profile of a *de novo* response in antigen naïve women ^47^. The relatively poorer transfer of resulting IgG3 subclass buts its greater and earlier induction following initial exposure as compared to well-transferred IgG1-biased responses common after recall is expected to be a factor in optimizing the timing of maternal immunization to optimally protect both mothers and neonates. Secondly, although SARS-CoV-2 mRNA vaccines elicit robust humoral immunity, accumulating evidence points to platform-specific class switch recombination (CSR). Unlike other vectors or COVID-19 vaccines ^48-51^, sequential mRNA dosing drives CSR toward IgG4, the terminal IgG subclass in the human Ig locus, and expands IgG4+ memory B-cell pools months after priming, with further amplification and elevation of antigen-specific IgG4 detected in serum after first and second boosters ^52-54^. Importantly this phenomenon has not been investigated during pregnancy, a setting in which it may be expected to have implications for vaccine efficacy given the more limited evidence of robust IgG4 transfer across the placenta ^55,56^, its functional monovalency due to Fab arm exchange ^57^, relatively poor effector function ^58^, and because repeated vaccination might be recommended during successive pregnancies and drive further shifts.

Thus, to complement the established literature and refine existing paradigms, we employed a comprehensive systems serology approach to characterize antibody responses over time and assess transplacental antibody transfer in pregnant individuals, as compared with responses observed in non-pregnant female adults immunized with BNT162b2 mRNA vaccine. Beyond systems serology, we proteomically identified SARS-CoV-2 vaccine-specific clonotypes that were detected as IgG4 in serum approximately six months after immunization to provide insights into the origin and histories of antibody clonotypes that class switch to IgG4. Resulting insights into the impact of pregnancy on responses to an unfamiliar antigen and in the context of novel mRNA vectoring strategies suggest effective strategies for protecting mothers and their newborns at the same time as defining new landscape in humoral immunity.

## Results

### Pregnancy alters vaccine-induced antibody phenotypes

We prospectively enrolled non-pregnant (n=65) and pregnant (n=36) female individuals receiving the BNT162b2 mRNA vaccine and collected serum samples at baseline (day 0), at the time of second primary vaccine dose (21-35 days later), and seven- and 28-days post second primary vaccine dose. Maternal and cord blood serum samples were obtained at the time of delivery, approximately six months post second primary vaccine dose. Serum was also collected at the time of a third dose for a subset of non-pregnant study participants and 159 days post second primary vaccine dose in pregnant participants. Non-pregnant individuals (median age = 42) were somewhat older than pregnant participants (median age = 31), and the non-pregnant cohort included one male participant. (**Supplemental Table 1**). Samples were collected 2021-22 in Belgium, when the predominant circulating strains were the Alpha and Beta variants of SARS-CoV-2 ^59^. Eight pregnant and four non-pregnant participants presented with hybrid immunity, as defined by seropositivity to SARS-CoV-2 nucleoprotein (N) at baseline or by a previous positive molecular SARS-CoV-2 PCR test at the beginning of the study, and were excluded from analysis except when specifically noted.

To define the potential impact of pregnancy on humoral immune responses, the magnitude, specificity, and Fc characteristics of antigen-specific antibodies induced by mRNA immunization were assessed. To delineate these changes, we employed systems serology approaches to measure antibody responses to recombinant whole spike, spike subunits S1, S2, and the receptor binding domain (RBD), and proline stabilized spike proteins from selected endemic, pandemic, and variant of concern (VOC) strains (**Supplemental Table 2**). Whereas unsupervised dimensionality reduction of antibody profiles indicated that both groups were similar at baseline (**Supplemental Figure 1A**), subsequent timepoints showed evidence of distinction (**Figure 1A**): seven days post second dose, serum antibody profiles of pregnant versus non-pregnant participants defined by Uniform Manifold Approximation (UMAP) showed differences between groups that began to be apparent as early as the day of second dose and were maintained at later timepoints (**Supplemental Figure 1B,C**). For a subset of participants, virus neutralizing activity in serum was evaluated at baseline and 28 days post second primary vaccine dose (**Figure 1B**). This activity was similar in pregnant and non-pregnant participants (p = 0.5956 by two-tailed Mann Whitney t test). In contrast, responses to S2P (prefusion-stabilized spike) vaccine antigen were robust in both groups by the time of and following the second vaccine dose (**Figure 1C**). Vaccine-specific Ig of each isotype evaluated (IgM, IgA, and IgG) showed a small but statistically significant elevation in non-pregnant as compared to pregnant participants at at least one timepoint.

**Fig 1.**
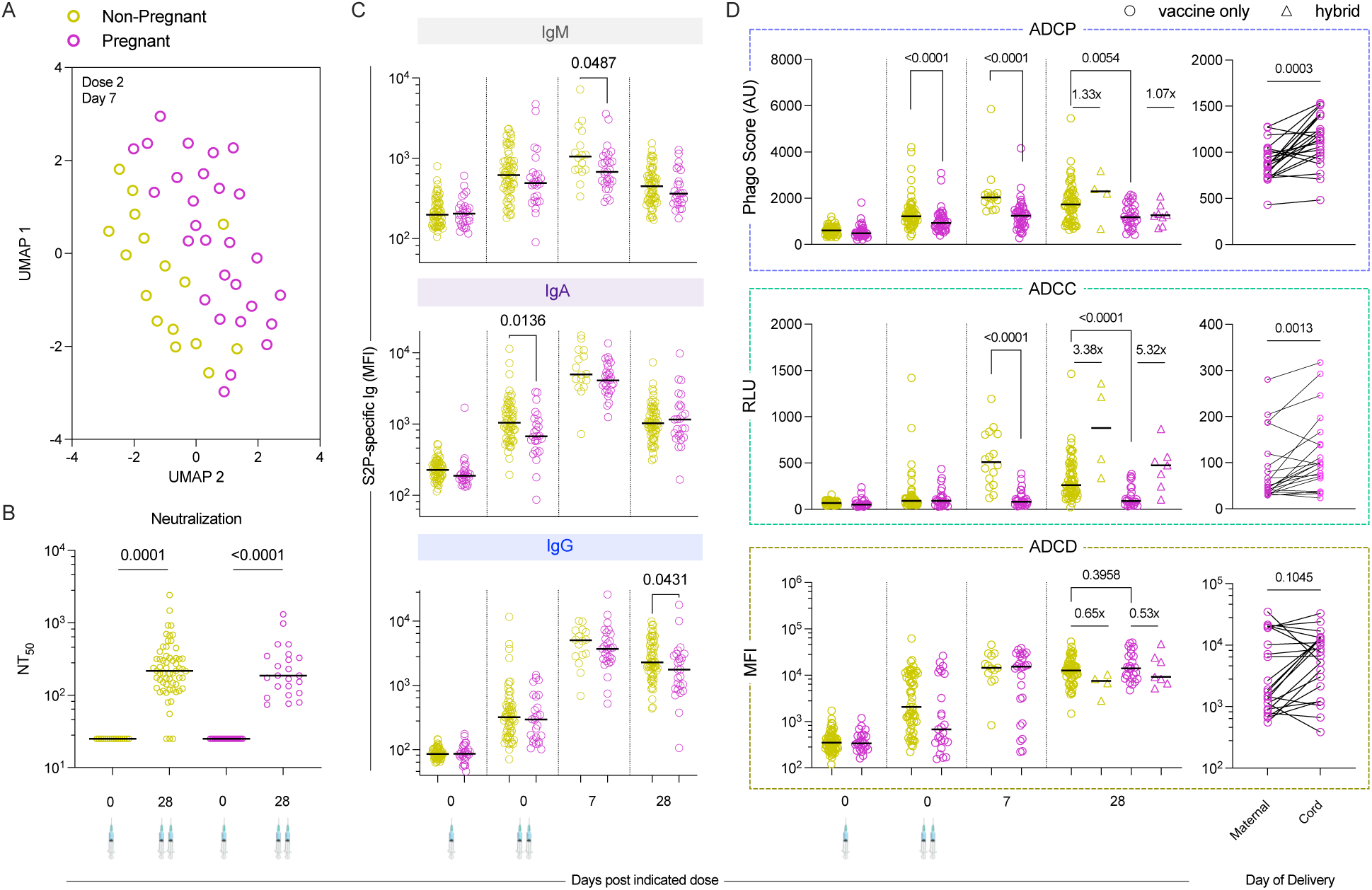
Phenotypes and functions of SARS-CoV-2-specific antibodies in pregnant (n=36) and non-pregnant (n=65) individuals following vaccination with BNT162b2. **A.** UMAP plot showing vaccine antigen-specific response features after dimensionality reduction in non-pregnant (green, n=28) and pregnant (pink, n=16) individuals at seven days post second dose. **B.** Neutralization titers (NT50) in serum samples from vaccinated non-pregnant and pregnant individuals at baseline and twenty-eight days post second dose. **C.** Median Fluorescent Intensity (MFI) of IgM (top), IgA (center), and IgG (bottom) responses to proline-stabilized spike (S2P) of SARS-CoV-2 as defined by multiplex assay over time. **D.** Antibody-effector functional activity directed to S2P in vaccinated pregnant and non-pregnant individuals (left) over time and for cord blood and maternal serum obtained at the time of delivery (right); Antibody-dependent Phagocytosis (top), Cellular Cytotoxicity (middle), and Complement deposition (Bottom) were measured. Bars indicate median fold changes between vaccinated individuals and those with hybrid immunity (infected and vaccinated). Statistical significance was assessed by the Mann–Whitney test for group differences and Wilcoxon rank sum test for paired maternal and cord blood samples. Functional activity is reported in arbitrary units (AU), relative light units (RLU), and median fluorescent intensity (MFI). Samples from all participants were not available for all timepoints.

### Vaccine-induced antibody effector functions

The potential functional consequences of these distinct response kinetics and antibody profiles were next evaluated using assays of antibody effector function against vaccine antigen, including phagocytosis, FcγRIIIa signaling as a surrogate for ADCC, and complement cascade factor C3b deposition. Higher levels of some but not all of these effector functions were observed at one or more timepoints in non-pregnant as compared to pregnant participants (**Figure 1D**). Whereas antibodies promoting phagocytosis were elevated in non-pregnant participants as early as the time of the second dose, these differences were not observed until seven days later for FcγRIIIa signaling. Differences in activity persisted until the 28-day post-second dose timepoint and were observed despite the induction of activating IgG subclasses and antibodies bound by FcγR (**Supplemental Figure 1D**).

### Potentiated effector function activity in the context of hybrid immunity

Although the number of participants was limited, several individuals initially enrolled in the study were subsequently found to exhibit nucleocapsid-specific antibody responses at baseline. While these participants were excluded from analyses assessing the impacts of pregnancy status, they provided an opportunity to evaluate the functional implications of hybrid immunity in the context of pregnancy. Because of the small number of individuals with this evidence of prior infection at baseline, formal statistical analyses were not performed. Nonetheless, ADCC activity of serum antibodies induced by the prior infection followed by vaccination appeared to be superior to that induced by vaccination alone, independent of pregnancy status (**Figure 1D**). In contrast, ADCP and ADCD responses were more comparable (within two-fold) between both groups.

### Efficient placental transfer of functional antibodies

Relevant to the intergenerational value of maternal immunization, we next investigated the transfer of vaccine-induced SARS-CoV-2-specific antibodies by comparing antibody responses and functional activity in maternal blood collected at the time of delivery with those in umbilical cord blood obtained from the fetal circulation. We observed significantly higher ADCC and ADCP activity (p=0.0013, p=0.0003 by Wilcoxon matched pairs signed rank test) in cord blood compared to maternal serum obtained at the time of parturition (**Figure 1D**). ADCD activity was not consistently elevated in cord blood but was robustly detected, with some dyads showing marked increases and others exhibiting either consistently high or low activity in both maternal and cord blood samples.

As expected, IgA and IgM antibodies, which are not transported across the placenta, were robustly elevated in maternal as compared to cord blood (**Supplemental Figure 2A).** Differences in IgG concentrations were more subtle but were typically greater in cord blood than in maternal blood, a phenomenon observed across both pandemic and endemic CoV (**Supplemental Figure 2B**). While there was variation among dyads, the median elevation of total CoV spike-specific IgG in cord relative to maternal blood was approximately two-fold. Among IgG subclasses, IgG1 exhibited the most efficient transfer, IgG2 transfer was variable, IgG3 levels were similar, and IgG4 levels were typically reduced in cord compared to maternal blood for SARS-CoV-2 antigen specificities (**Supplemental Figure 2B**). Antibodies to endemic CoV antigens exhibited generally similar transfer profiles, except for more efficient transfer of IgG4, resulting in similar levels in maternal and cord blood, in contrast to the generally lower levels observed for pandemic strains (**Supplemental Figure 2B**). For comparison, we also investigated transfer of tetanus toxoid-, diphtheria toxoid-, and pertactin-specific IgG to the fetus (**Supplemental Figure 2C**), as mothers received the Tdap vaccine in late second or early third trimester. Transfer ratio profiles similar to those for CoV-specific antibodies were observed across these vaccine antigens. Total IgG levels exhibited a median nearly two-fold elevation in cord blood, driven largely by preferential transfer of maternal IgG1 (**Supplemental Figure 2C**). Unlike IgG4 antibodies to SARS-CoV-2 and its VOC, but similar to endemic CoV antigens, no evidence of bias against the transfer of IgG4 was observed for these other vaccine antigens. Overall, in conjunction with increased levels of IgG1 and total S2P-specific IgG in cord blood, the elevated antibody functional activity observed in the cord blood samples may contribute to protection of infants during the initial months of life while their immune system is still maturing.

### Late induction of spike-specific IgG4

Several studies have reported continued evolution and class switching toward distal subclasses of IgG following repeated mRNA vaccine immunization ^48-54^. Samples collected from this cohort allowed longitudinal tracking of IgG subclass profiles of vaccine-specific antibodies over time in the context of only two vaccine doses and during pregnancy. Whereas robust increases in S2P-specific IgG1 and IgG3 levels were observed, responses to IgG2 and IgG4 were negligible up to 28 days post second dose (**Figure 2A**). However, levels of these inflammatory IgG subclasses dropped by more than an order of magnitude by 5 months post second dose, a timepoint at which IgG2 responses stayed low, but IgG4 responses became elevated (**Figure 2A**). Late elevation of vaccine-specific IgG4 was observed in both pregnant and in the subset of non-pregnant participants who returned to receive a third dose of vaccine and for whom a comparable timepoint was available. In the latter group, increasing IgG4 responses generalized across different spike domains and conformations but were not detected against nucleocapsid (**Figure 2B**), and appeared less pronounced in participants with hybrid immunity (**Supplemental Figure 3**). IgG4 responses cross-reacted with alpha, beta, gamma, delta, kappa, mu, SARS-CoV-1, MERS, OC43, and HKU1 spike proteins, although responses to the endemic strains were quite low (**Figure 2C**). In contrast, elevated IgG4 to control antigens were not observed (**Figure 2C**). Overall, over the period during which total S2P-specific IgG declined seven-fold (**Figure 2D**), the relative composition of this antibody pool, assessed by median fluorescent intensity of signals for each subclass, changed markedly (**Figure 2E**).

**Fig 2.**
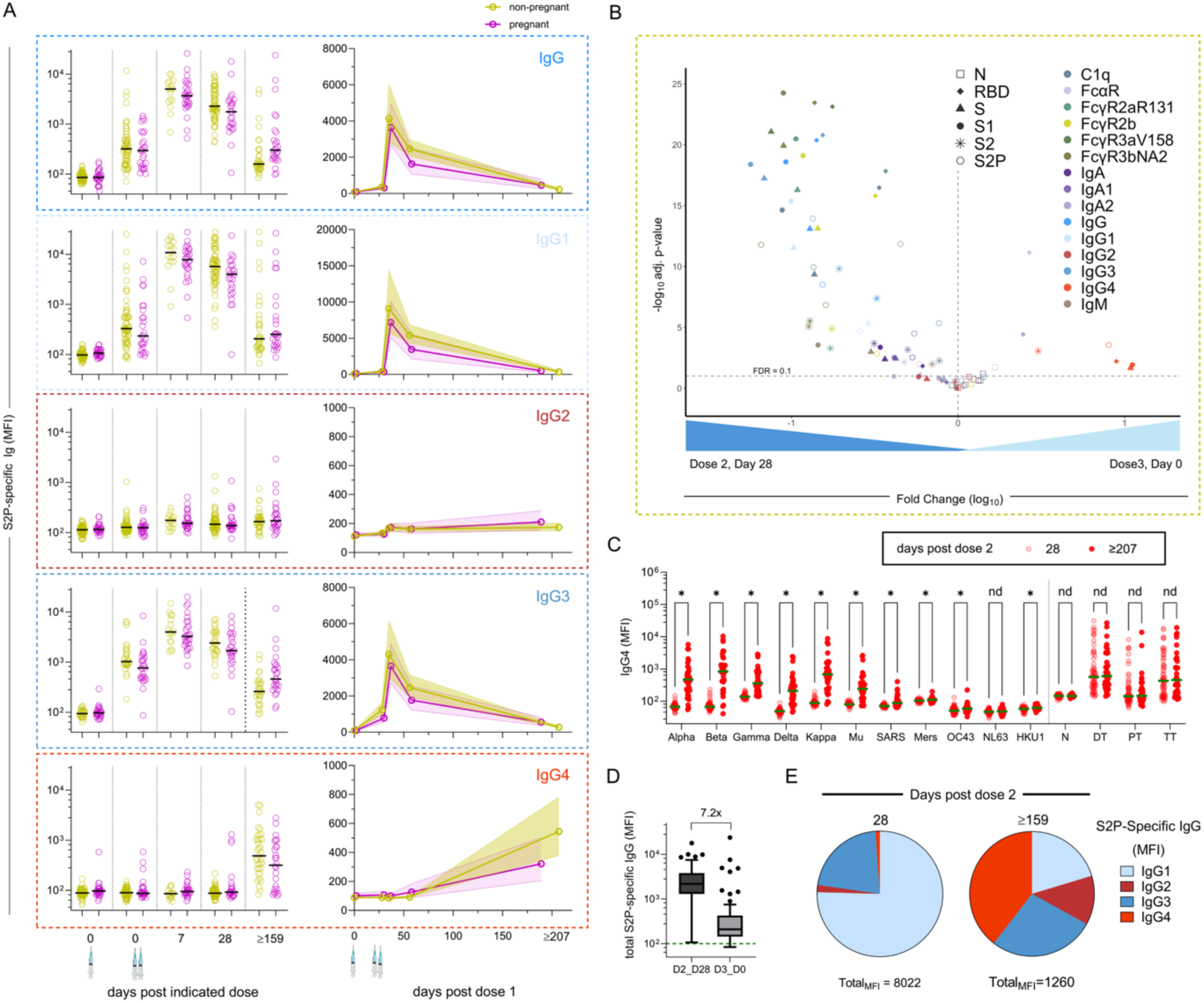
COVID-specific IgG4 class switching increases over time after only two doses of BNT162b2. **A.** (Left) Median Fluorescent Intensity (MFI) of total IgG and each IgG subclass specific to S2P as defined by multiplex assay for non-pregnant (green) and pregnant (pink) individuals over time. Bar indicates the median response. (Right) Line-graph showing geometric mean in solid lines for non-pregnant and pregnant individuals and 95% confidence interval defined by the shaded regions. **B.** Volcano plot showing the fold change (x-axis) and statistical significance (Mann Whitney test with Benjamini Hochberg correction, y-axis) of differences in levels of each antibody type between twenty-eight days post dose two and at the time of third dose in non-pregnant individuals. Shapes indicate antigen, color indicates antibody isotype and binding to Fc receptors. **C.** Levels of IgG4 detected across coronavirus and control antigens (DT - Diphtheria toxoid, PT – pertactin, TT – Tetanus toxoid) at twenty-eight days post second dose (light pink) and ∼210 days later at the time of third dose (red). Statistical significance was determined by multiple T-tests with Benjamini-Hochberg correction and Q= 0.05. **D.** Boxplot of total S2P-specific IgG at twenty-eight days post second dose and at the time of third dose of mRNA vaccine in non-pregnant individuals. Bar denotes fold change in group medians. **E.** Pie-chart showing relative levels (MFI) of each IgG subclass at day twenty-eight (left) and six months (right) post second dose in pregnant individuals. Arithmetic sum of MFI for all IgG subclasses indicated at the bottom.

### Functional implications of late IgG4 responses

IgG4, which exhibits weaker effector functions than the IgG1 and IgG3 subclasses, has been variably observed to either contribute to or interfere with antiviral antibody activity, depending on factors such as antibody levels and antigen specificity ^60^. In the present study, statistically significant correlative relationships, either direct or indirect, between IgG4 levels and antibody functions were generally not observed (**Supplemental Figure 4**), suggesting that in these antibody pools, IgG4 does not make a strong contribution either favorably or unfavorably to host protection. Among effector functions, IgG4 has been reported to contribute most to phagocytic activity ^50,61^, but depletion of IgG4 from a subset of samples (**Supplemental Figure 5**) did not result in statistically significant changes in ADCP activity (**Supplemental Figure 6**).

### Features of secreted antibody clonotypes detected as IgG4

CSR to IgG4 could occur directly in naïve B cells or via sequential switching from upstream subclasses over time within a clonal lineage. As part of a separate study, spike-specific total IgG was isolated from sera collected from six pregnant participants in the cohort at the time of second primary vaccine dose (D2D0), 28 days after the second primary dose (D2D28), and at delivery (maternal and cord blood). These secreted antibodies were characterized at clonotype resolution by BCR-Seq and Ig-Seq to identify the sequence characteristics, diversity, and relative levels of spike-specific IgG secreted systemically ^62^. To identify which spike-specific IgG clonotypes had undergone class switching to IgG4, total IgG4 was affinity purified from sera collected at 159 days after the second primary dose and analyzed by mass spectrometry. Clonotypes identified in the purified IgG4 fraction were compared with previously defined spike-specific IgG repertoire to determine which antigen-specific clonotypes were detected as IgG4 at this late time point.

Across donors, up to four (mean = 1.8) distinct spike-specific clonotypes were detected as IgG4 among 12-145 (mean = 71.3) total spike-specific clonotypes in each donor, representing 0-8.3% of clonotypes by abundance (mean = 3.96%) (**Figure 3A**). The relative abundance of clonotypes detected as IgG4 could be tracked over time in maternal serum and cord blood. Notably, 91% (10 of 11) of the clonotypes detected as IgG4 at the late timepoint were observed to be present at all sampled time points (**Figure 3B**), a feature observed for only 20.3% of spike-specific clonotypes. Clonotypes detected as IgG4 also tended to be among the more abundant and more highly somatically hypermutated clonotypes. They generally exceeded the median abundance of clonotypes not detected as IgG4 within the same donor and time point (**Figure 3B**) and showed higher median levels of somatic hypermutation (**Figure 3C**). Consistent with longitudinal trends in serum, clonotypes detected as IgG4 increased in relative abundance from 28 days after the second primary dose to delivery (**Figure 3D,E**). In agreement with the relatively poor placental transfer of IgG4 (**Supplemental Figure 2B**), these clonotypes, particularly those that were more abundant, were detected at modestly lower levels in cord blood than in maternal blood at delivery (**Figure 3F**).

**Fig 3.**
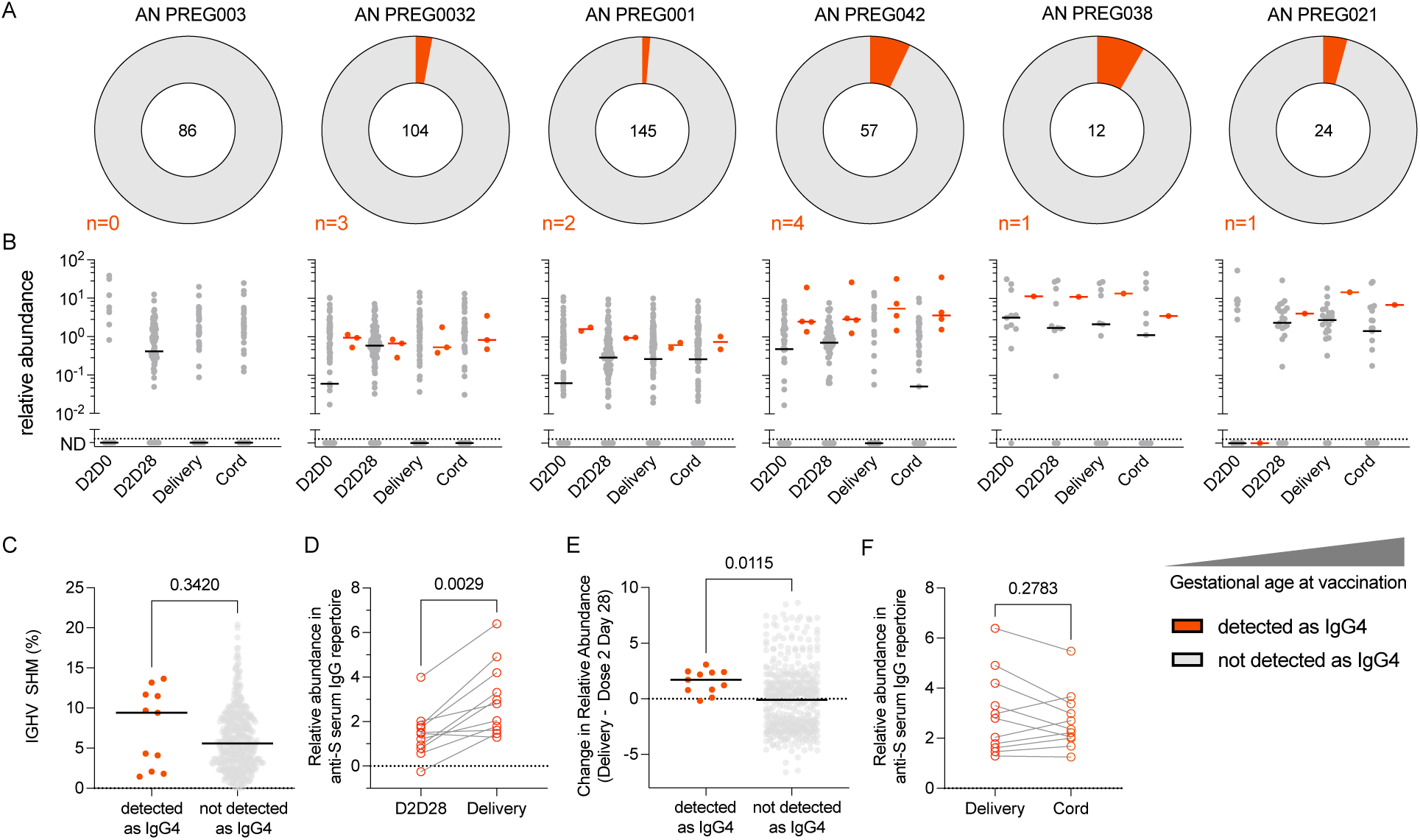
Vaccine-elicited SARS-CoV-2 spike-specific IgG clonotypes are later detected as IgG4. A. Donut charts depicting proportion and number of spike-specific IgG clonotypes detected as IgG4 (red) versus those not detected as IgG4 (gray) for each of six donors arranged in order of increasing gestational age at time of vaccination. The total number of spike-specific IgG clonotypes is shown at the center of each chart, and the number of clonotypes detected as IgG4 is indicated below in red. **B.** Relative abundance of each spike-specific clonotype detected as IgG4 (red) versus clonotypes not detected as IgG4 (gray) over time. Horizontal bars denote the median. **C.** Somatic hypermutation (SHM) levels for each serum clonotype detected as IgG4 (red) versus clonotypes not detected as IgG4 (gray). **D.** Relative abundance of clonotypes detected as IgG4 at 28 days post second dose (D2D28) and at delivery. **E.** Change in relative abundance of clonotypes detected as IgG4 (red) versus clonotypes not detected as IgG4 (gray) from D2D28 to delivery. **F.** Relative abundance of IgG4-associated clonotypes in maternal versus cord blood samples.

## Discussion

Pregnancy is a unique physiological state in which immune adaptations are essential for safeguarding both maternal and neonatal health while ensuring maternal tolerance towards a semi-allogenic fetus. Previous studies of responses to vaccination have shown variable impacts of pregnancy status: while some studies have reported reduced immunogenicity ^63,64^, others have suggested comparable humoral immune responses ^35,47,65,66^. Much of the SARS-CoV-2 vaccine literature was generated during the height of the pandemic, often relying on convenience cohorts with heterogeneous sampling schedules and limited longitudinal follow-up. In contrast, this study benefitted from prospective and simultaneous enrollment of pregnant and non-pregnant women from a single geographic area. Longitudinal sampling enabled a more detailed understanding of vaccine-induced kinetics during pregnancy as well as antibody transfer to the developing fetus.

Our findings revealed that while binding antibody responses were relatively comparable between pregnant and non-pregnant women, ADCP and ADCC were diminished in the pregnant cohort. This functional adaptation despite more quantitatively similar antibody levels suggests that pregnancy may induce qualitative modifications to circulating immunoglobulins but has not been consistently observed in other studies ^35,47,63-66^, and may therefore result from other confounding factors, such as differences in donor age, which was greater in non-pregnant participants. Given the well-established efficacy of SARS-CoV-2 immunization in protecting pregnant women and infants from COVID-19 morbidity and mortality ^67-71^, these differences may be of limited clinical relevance.

Concurrently, the conservation of late IgG4 class-switching in pregnant women indicates that subclass redistribution dynamics following repeated mRNA vaccination are not substantially modified by pregnancy-associated immune adaptations. This profile has been observed across the lifespan, including in children and the elderly ^72,73^ and for both Moderna and Pfizer/BioNTech mRNA vaccines, but not with other vaccine platforms or after breakthrough infection ^74^. It has also been shown to persist in the context of antigenic updates in vaccines targeting VOC ^75^. Despite the continued overall efficacy of these vaccines ^76,77^, the magnitude of late CSR to IgG4 has also been reported to associate with increased relative risk of breakthrough infection ^78^. Overall, while increases in IgG4 after repeated SARS-CoV-2 mRNA vaccination are well documented, their clinical and functional consequences remain incompletely defined.

Though IgG4 is an antibody subclass distinguished by its relatively weak binding to activating Fc receptors, poorer complement activation, and more limited capacity to mediate pro-inflammatory effector responses ^79^, prior studies have shown that IgG4 can have both positive ^80,81^ and negative ^60^ effects on antibody effector function; our data are consistent with a limited functional impact. High inter-individual variability within complex polyclonal antibody pools likely obscures the potential influence of switching to this less inflammatory subclass in our relatively small cohort. In prior work, addition of IgG4 monoclonals reduced the antiviral activity of IgG1 monoclonals of the same epitope specificity but failed to drive similar reductions in the activity of polyclonal antibody pools ^82^. Nonetheless, relationships between elevated IgG4 and decreased effector function have sometimes, but not always ^83^, been observed in other studies of polyclonal serum responses directed at SARS-CoV-2 ^73^ or other viruses ^60,84^. Consistent with this apparent inconsistency, both modeling and experiments to the define the effect of IgG4 on effector functions mediated better by IgG1 and IgG3 subclasses demonstrate a bidirectional effect depending on the composition of the overall antibody pool and the magnitude of responses ^50^. The resulting model suggests that at high IgG titers, increased IgG4 acts primarily as a competitive inhibitor, blocking proinflammatory subclasses from binding antigen and resulting in reduced binding to FcγR, By contrast, when total antigen-specific antibody levels are low, IgG4 can lead to increased effector function, as its presence increases overall amount of antigen-bound antibody, enhancing FcγR-mediated functions ^50^. In the present study, we only directly tested the impact of the IgG4 pool on ADCP activity. While depletion of IgG4 increased phagocytic activity in some donors, it decreased it in others. Overall, the relevance of antibody effector functions to providing protection from disease, especially in the context of continued antigenic variation and escape from neutralizing activity motivates continued study of the impact of late CSR to IgG4 ^85-87^.

BCR-Seq and Ig-Seq analyses uncovered several novel observations regarding IgG4 class-switching following mRNA vaccination that build upon and extend recent studies documenting this phenomenon ^52,53,88^ by demonstrating previously unknown attributes of IgG4-switched serum clonotypes. Although our data do not exclude the possibility that the late IgG4 response also includes *de novo* clonotypes induced from naïve B cells only at late time points, prior studies of SARS-CoV-2 recall responses indicate that secondary serum antibody responses are dominated by pre-existing clonotypes established during earlier exposures ^89^. Here, longitudinal clonotype tracking revealed that most clonotypes detected as IgG4 at the late timepoint were present at all earlier timepoints. This observation suggests sequential CSR among established, antigen-experienced clonotypes occurs over time. Clonotypes detected as IgG4 tended to be more prevalent, more persistent, and more somatically hypermutated than clonotypes not detected as IgG4, suggesting that switching is associated with clonotype size and degree of stimulation. Future studies examining the balance between *de novo* and sequential class-switching, the B cell subsets and anatomical compartments involved, and the signals that orchestrate this process will enhance our understanding of human humoral immunity and may inform efforts to optimize vaccine design.

Many questions remain. SARS-CoV-2 mRNA platforms appear particularly prone to inducing late class switching toward distal subclasses, possibly due to prolonged antigen availability and germinal center reactions ^90^. Similar IgG4 responses have been seen after repeated vaccination in the context of protein-based DTaP and HIV vaccines ^60,91^. Repetitive exposure to allergens such as bee venom, peanuts, eggs, and milk, including intentional exposure as a part of tolerization therapies, are also associated with CSR to IgG4 ^92-95^. While the constant region to which a B cell switches is modulated by cytokines and B cell activators at the level of transcription of non-rearranged heavy-chain constant genes ^96^, the regulators for germline transcription of the γ4 gene locus are not very well understood in humans ^52^, though Interleukin-4 (IL-4) in tandem with IL-10 ^97^ and tolerance-promoting signals from regulatory B and T cells ^98^ have been described to be involved in switching to IgG4. Whether late IgG4 class-switching reflects a physiological mechanism to dampen prolonged immune activation or has implications for durability and degree of vaccine-induced protection remains to be fully understood, but may be important in the context of potential seasonal or pregnancy-associated booster doses of for combination mRNA vaccines for influenza and COVID, Limitations exist. Longitudinal long-read sequencing would be needed to definitively CSR histories across individual clonotypes. Additionally, in-depth clonotype analysis was performed for only a subset of participants, and the extent of IgG4 switching varied among individuals; analysis of additional vaccine recipients will be needed to confirm and extend these observations, including in non-pregnant vaccine recipients. Neutralization tests were performed on a subset of samples and timepoints, and assays of effector function were conducted with recombinant antigen and cell lines as opposed to more biologically-relevant targets and effectors. Other limitations include unequal and relatively small group sizes, compounded by some missing samples, which affects the statistical power and comparability of group-level analyses. Moreover, non-pregnant participants were generally older, potentially introducing residual confounding given the known influence of age on immune function and antibody responses. These factors should be considered when interpreting the findings.

In summary, this work observed adaptations of the humoral immune response to immunization associated with pregnancy that included somewhat lower vaccine-specific antibody levels and Fc effector function, an observation that has been made for this and other vaccines that remain effective in protecting mothers and infants. However, both pregnant and non-pregnant vaccine recipients exhibited CSR toward the more functionally inert IgG4 subclass late after the second vaccine dose. The clonotypes detected as IgG4 at late timepoints tended to be more highly mutated, more prevalent, and to increase over time. Elucidating the balance of IgG4 antibodies that arise from *de novo* class-switched plasmablasts versus serial CSR, and further research into the specific B cell subsets, germinal center or extrafollicular dynamics, and signals that drive late CSR to IgG4 promise to clarify our understanding of humoral responses in humans and may help optimize future vaccine development,.

## Methods

### Ethics Statement

All study participants provided written informed consent. The study protocols were reviewed and approved by the Ethics Committee (EC) of the Institute of Tropical Medicine (ITM), Antwerp, Belgium; the EC of the Erasme Hospital, Brussels, Belgium; the Local EC of the University Hospital Center of St Pierre, Brussels, Belgium; the EC of the Antwerp University Hospital, Antwerp, Belgium; the Belgian Federal Agency for Medicines and Health Products, and Dartmouth College.

### Human Subjects

Pregnant (N=36) and non-pregnant (N=65) participants were recruited in three different clinical trials conducted in Belgium (PregCoVac, NCT05618548; PICOV-VAC, NCT04527614; REDU-VAC, NCT04852861) (**Supplemental Table 1**). Participants were asked about their previous SARS-CoV-2 exposure. Alpha, Beta and Delta variants were the dominant strains at the time of sample collection, however infecting viral strains were not characterized or genotyped. All participants were vaccinated with the primary series of two doses of BNT162b2 mRNA COVID-19 vaccine, and a subset of non-pregnant participants received a third, booster, dose. Serum was prepared from blood collected at baseline (time of first vaccine dose), at the time of the second dose (21-35 days after the first dose), and at seven and twenty-eight-days after the second dose for both groups. For non-pregnant participants, samples were additionally collected at the time of the third dose when available. For pregnant participants, serum samples were also collected approximately 159 days after the second vaccine dose. Both maternal and cord blood samples were collected at the time of delivery for pregnant participants (8-217 days after the initial dose). Participants were considered previously infected if they met any of the following criteria (a) a prior positive molecular SARS-CoV-2 PCR test, (b) detectable SARS-CoV-2 nucleocapsid-specific antibodies at any tested time point before administration of a third vaccine dose, or (c) an increase in titer of SARS-CoV-2 receptor binding domain (RBD) antibodies between available time points not encompassing a vaccination event, as previously described ^99^.

### Fc Array

Serum antibodies specific for a panel of COVID-specific antigens were sourced commercially or made in house using Expi293 or HEK293 cells and purified via affinity chromatography (**Supplemental Table 2**) and characterized using the Fc multiplexed assay as described previously ^100^. Briefly, antigens were covalently conjugated to fluorescently coded magnetic beads (MagPlex Microspheres, Luminex Corporation). Serum dilutions appropriate for each detection reagent, varying from 1:500 to 1:5000, were prepared in Assay Wash Buffer (AWB) (Luminex Corporation) to optimize assay performance based on pilot testing and prior experience, added to washed beads, and incubated for 2 hours. Following binding and washing, antigen-specific antibodies were detected by R-phycoerythrin (PE)-conjugated secondary reagents specific to human immunoglobulin isotypes and subclasses or site-specifically tetramerized Fc receptors (**Supplemental Table 2**). Following standard incubation and washing processes, beads were resuspended in AWB, and Median Fluorescent Intensity (MFI) values were acquired on a FlexMap 3D array reader (Luminex Corporation). A known SARS-CoV-2 seropositive serum sample and the monoclonal antibody S309 ^101^ that binds to RBD of the spike glycoprotein were used as positive controls; a known SARS-CoV-2 seronegative sample was diluted in PBS-TBN (0.1% BSA, 0.02% sodium azide, 150 mM sodium chloride, 50 mM sodium phosphate monobasic anhydrous, 0.05% Tween-20, pH 7.4) buffer and buffer alone blanks served as biological and technical negative controls.

### Neutralization

Neutralizing antibodies (nAb) against SARS-CoV-2 were quantified as described previously ^102^ for a subset of enrolled participants. Heat-inactivated serum was serially diluted (1:50 to 1:25,600) in EMEM containing 2 mM L-glutamine, 100 U/mL–100 µg/mL penicillin–streptomycin, and 2% fetal bovine serum, then incubated for 1 hour at 37°C and 7% CO_2_ with 3×TCID_100_ of Wuhan strain (2019-nCoV-Italy-INMI1, 008 V-03893). After incubation, 100 µL of the virus–serum mixture was added to 100 µL of Vero cells (18,000 cells/well) in 96-well plates and cultured for 5 days at 37°C and 7% CO_2_. Viral cytopathic effects were assessed by microscopy, and the Reed–Muench method was used to determine the neutralizing titer that reduced infection by 50% (NT_50_), which served as the nAb measure. Each assay run included an internal reference standard (a pooled serum from naturally infected and vaccinated adults) that was calibrated to the WHO International Standard 21/234 (NIBSC).

### Reporter cell assay of antibody-dependent cellular cytotoxicity (ADCC)

A surrogate for ADCC activity was measured using a CD16^+^ (FcγRIIIa) reporter assay system. Jurkat Lucia NFAT (Invivogen, jktl-nfat-cd16) cells were cultured according to the manufacturer’s instructions. Cultured cells express CD16, which, when engaged on the cell surface, leads to luciferase expression and secretion. First, high-binding 96-well plates were coated overnight at 4 °C with 1 μg/mL of S2P antigen ^103^. Following incubation, plates were washed (PBS + 0.1% Tween 20) and blocked (PBS + 2.5% BSA) at room temperature (RT) for 1 h. Following plate washing, 100,000 cells per well and 1:100 dilution of serum samples were added to each well in cell culture media lacking antibiotics in a 200 μL volume. Following 24 h incubation, 25 μL of supernatant from each well was transferred into a white optiplex (Fisher Scientific cat # 50-209-9794), 96-well plate, and after adding 75 μL of Quantiluc substrate (Invivogen cat # rep-qlc4lg1), the plates were read immediately on a Spectramax luminometer (Molecular Devices) using 1s integration time. Kinetic readings at 0 min, 2.5 min and 5 min were measured, and the mean reading was noted. The HIV-specific monoclonal antibody VRC01 ^104^ was used as a negative control; the cell stimulation cocktail (Thermo Fischer Scientific, 00-4970-93) and SARS-CoV-2-specific monoclonal antibody S309 served as positive controls. Serial dilution of a known seropositive sample was used to set the concentration of the serum for the assay.

### Antibody-dependent complement deposition (ADCD)

ADCD experiments were performed as previously described ^105^. Briefly, serum samples were first heat-inactivated by incubation at 56°C for 30 min. Samples were then incubated on a shaker for 2 h at RT with antigen-conjugated multiplex assay beads as described above. The optimal dilutions for serum were determined from a pilot experiment that tested a subset of samples over a range of dilutions to define the linear ranges and 1:2000 dilution of serum was used in the assay. Guinea pig complement serum (Cedarlane # CL4051) was diluted 1:60 in gelatin veronal buffer (Sigma-Aldrich, GVB++, G6514) and mixed with dilute serum samples at 37 °C with shaking for 20 min. Following this incubation, cold EDTA solution (15 mM in 1X PBS) was added to prevent further complement activation. After washing, samples were incubated with biotin-conjugated goat anti-C3b (1 µg/mL) (Icllab GC3-60B-Z) at room temperature for 1 h, followed by staining with Streptavidin-R-Phycoerythrin (Millipore Sigma 42250) reagent per the manufacturer’s recommendation. A final wash was performed, and samples were resuspended in Luminex sheath fluid and MFI acquired on a FlexMap 3D reader (Luminex). Assay controls included a technical blank (no antibody) heat-inactivated complement serum, S309 antibody, a known negative and positive serum sample.

### Antibody-dependent cellular phagocytosis (ADCP)

The phagocytic potential of the serum antibodies was assessed. Briefly, 1 μM neutravidin-labeled yellow-green, fluorescent beads (Thermo Fisher, F8776) were conjugated^106^ to biotinylated S2P antigen. Beads were subsequently spun down and washed twice in 1% BSA in PBS (PBSA) to remove excess unbound antigen and then resuspended at a final dilution of 1:100 in PBSA. Conjugated beads (0.1 µl of the supplied beads or 10 µl of the dilution above) were then incubated with serum samples for 2 hrs to facilitate the formation of Ag-Ab complexes. Following incubation, 25000 THP-1 cells/well (ATCC, TIB-202) were added to the complexes in total of 200 µl THP-1 medium (RPMI with 10% FBS+ 1x Pen-Strep)) and incubated or 16 hours at 37°C and 5% CO_2_. Cells were fixed and analyzed using flow cytometry (Novocyte-Agilent Technologies). Scores were calculated as the percentage of cells that phagocytosed one or more fluorescent beads multiplied by the MFI of this population. S309, which was used to adjust for plate-to-plate variation, and VRC01 antibodies were included as positive and negative controls, respectively. Additional control wells with no added antibody were used to determine the level of antibody-independent phagocytosis. An initial pilot experiment helped in determining the optimal serum concentration and signal-to-noise ratio and 1:100 dilution of serum was used to perform the experiment.

### IgG4 Depletion

IgG4 capture beads were made by reacting to 10 mg of M-270 Streptavidin Dynabeads (Invitrogen) with 100 μg of a biotin-conjugated IgG4-specific Ab (Thermo-Fisher). The beads were washed three times with 500 μL of PBS and then incubated with the Ab (in PBS) for 30 min at room temperature with gentle rotation. After conjugation, the beads were washed three times with 0.1% BSA in PBS and stored at a final concentration of 10 mg/mL. For IgG4 depletion, 6 μL of dose 2 day 159 participant sera was added to 50 μL of beads and incubated at 4 °C for 24 h. Beads were then removed via magnetic separation. Supernatant was collected was collected as the IgG4 depleted fraction. A Luminex-based subclassing assay was run before and after the bead incubation to assess the efficiency of IgG4 depletion. A selection of samples, before and after depletion, was also assessed for their remaining nondepleted IgG subclass profiles. Nonspecific IgG subclass loss was not observed.

### High-throughput sequencing of V_H_

Total RNA (500 ng) extracted from PBMC taken 7 days after vaccination with dose 2 was used for library preparation. Reverse transcription was performed using SuperScript IV (Invitrogen) and Oligo(dT) primer (Invitrogen) according to manufacturer instructions. V_H_-specific transcripts were amplified using the FastStart High Fidelity PCR System (Roche) with gene-specific primers2. Single-chain amplicons were barcoded using the Next® UltraTM II DNA Library Prep Kit (NEB, E7645) and sequenced on the Illumina NextSeq platform.

### LC-MS/MS Analysis

IgG4 was captured from participant sera (dose 2 day 159) using the same IgG4-specific streptavidin bead preparation and incubation conditions described above for IgG4 depletion. Following magnetic separation, bound IgG4 was subsequently eluted by three sequential incubations with 0.5 mL of 1% (v/v) formic acid. Elution fractions were pooled and concentrated under vacuum to a volume of ∼10 μL and neutralized using 2 M NaOH. The neutralized elution samples and flowthrough samples were denatured with 50 μL of 2,2,2-trifluoroethanol (TFE) and 5 μL of 100 mM dithiothreitol (DTT) at 55°C for 1 hr and then alkylated by incubation with 3 μL of 550 mM iodoacetamide for 30 min at RT in the dark. Alkylation was quenched with 892 μL of 40 mM Tris-HCl, and protein was digested with trypsin (1:30 (w/w) trypsin/protein) for 16 hr at 37°C. Formic acid was added to 1% (v/v) to quench the digestion, and the sample volume was concentrated to 150 μL under vacuum. Peptides were then purified using C18 spin columns (Thermo Scientific, 89870), washed three times with 0.1% formic acid, and eluted with a 60% acetonitrile and 0.1% formic acid solution. C18 eluate was concentrated under vacuum centrifugation and resuspended in 50 μL in 5% acetonitrile, 0.1% formic acid.

Samples were analyzed by liquid chromatography-tandem mass spectrometry (LC-MS/MS) on an Easy-nLC 1200 (Thermo Fisher Scientific) coupled to an Orbitrap Fusion Tribrid (Thermo Scientific). Peptides were first loaded onto an Acclaim PepMap RSLC NanoTrap column (Dionex; Thermo Scientific) prior to separation on a 75 μm × 15 cm Acclaim PepMap RSLC C18 column (Dionex; Thermo Scientific) using a 1.6%–76% (v/v) acetonitrile gradient over 90 mins at 300 nL/min. Eluting peptides were injected directly into the mass spectrometer using an EASY-Spray source (Thermo Scientific). The instrument was operated in data-dependent mode with parent ion scans (MS1) collected at 120,000 resolution. Monoisotopic precursor selection and charge state screening were enabled. Ions with charge ≥ +2 were selected for collision-induced dissociation fragmentation spectrum acquisition (MS2) in the ion trap, with a maximum of 20 MS2 scans per MS1. Dynamic exclusion was active with a 15-s exclusion time for ions selected more than twice in a 30-s window. Each sample was run three times to generate technical replicate datasets.

### IgG4 MS/MS Data Analysis

Donor-specific peptide search databases for MS data acquisition were created using V_H_ sequences. V_H_ amino acid sequences with ≥2 reads from high-throughput sequencing were included to construct the peptide search database. These V_H_ sequences were then combined with a database of background proteins, which included a consensus human protein database (Ensembl 73, longest sequence/gene), and a list of common protein contaminants (MaxQuant). Spectra were searched against the database using SEQUEST (Proteome Discoverer 2.4; Thermo Scientific). A precursor mass tolerance of 5 ppm and fragment mass tolerance of 0.5 Da were used. Modifications of carbamidomethyl cysteine (static), oxidized methionine, and formylated lysine, serine or threonine (dynamic) were selected. High-confidence peptide-spectrum matches (PSMs) were filtered at a false discovery rate of <1% as calculated by Percolator (q-value <0.01, Proteome Discoverer 2.4; Thermo Scientific). Iso/Leu sequence variants were collapsed into single peptide groups. For each scan, PSMs were ranked first by posterior error probability (PEP), then q-value, and finally cross-correlation (XCorr) score. Only unambiguous top-ranked PSMs were kept; scans with multiple top-ranked PSMs (equivalent PEP, q-value, and XCorr) were designated ambiguous identifications and removed. The average mass deviation (AMD) for each peptide was calculated as previously described^7^. Peptides with AMD >1.5 ppm were removed.

High-confidence peptides identified were then mapped to clonotype clusters. Peptides that are uniquely mapped to a single clonotype were considered ‘informative’, and clonotypes detected ≥2 PSM were kept for as high-confidence identifications. The abundance of each antibody clonotype was calculated by summing the extracted-ion chromatogram (XIC) areas of the informative peptides mapping to ≥4 amino acids of the CDRH3 region.

IgG4 serum clonotypes that were clonally related to spike-specific serum IgG clonotypes were classified as detected as IgG4 if an identical CDRH3 peptide in the IgG4 MS dataset was present among the spike-specific IgG clonotypes and was detected in each of the triplicate IgG4 MS injections. To ensure confident assignment, clonotypes with low signal, defined as less than 0.1% relative abundance at the delivery time point, were excluded.

### Statistical Analysis

Statistical and graphical analysis was performed in GraphPad Prism (version 10.6.1.) and R Studio version 4.5.2 (Open Source).

## Author contributions

D.N.K., T.Y., N.M.T., and A.Y. designed and executed the experiments and analyzed the data. D.N.K., T.Y., N.K., J.A.W., and M.E.A. performed bioinformatic analysis and visualization. M.E.A., J.L., and A.M. conceived the project and supervised laboratory analyses. K.K.A. provided neutralization titer data. G.K., and A.H. provided technical support. A.M. and M.E.G. served as a principal clinical investigators for this study. L.D.B., S. D., A. V., N. D., and K.M. contributed to donor enrollment and clinical sample collection and management. P.P. helped with the management and logistics of the clinical samples. D.N.K., and M.E.A. wrote the manuscript with input from co-authors. All authors reviewed and edited the manuscript.

